# Tuning Strand Displacement Kinetics Enables Programmable ZTP Riboswitch Dynamic Range *in vivo*

**DOI:** 10.1101/2022.10.20.513036

**Authors:** David Z. Bushhouse, Julius B. Lucks

## Abstract

Recent work has shown that transcriptional riboswitches function through internal strand displacement mechanisms that guide the formation of alternative structures which drive regulatory outcomes. Here we sought to investigate this phenomenon using the *Clostridium beijerinckii pfl* ZTP riboswitch as a model system. Using functional mutagenesis with *in vivo* gene expression assays in *E. coli*, we show that mutations designed to slow strand displacement of the expression platform enable precise tuning of riboswitch dynamic range (2.4–34-fold), depending on the type of kinetic barrier introduced, and the position of the barrier relative to the strand displacement nucleation site. We also show that expression platforms from a range of different *Clostridium* ZTP riboswitches contain sequences that impose these barriers to affect dynamic range in these different contexts. Finally, we use sequence design to flip the regulatory logic of the riboswitch to create a transcriptional OFF-switch, and show that the same barriers to strand displacement tune dynamic range in this synthetic context. Together, our findings further elucidate how strand displacement can be manipulated to alter the riboswitch decision landscape, suggesting that this could be a mechanism by which evolution tunes riboswitch sequence, and providing an approach to optimize synthetic riboswitches for biotechnology applications.

## INTRODUCTION

Riboswitches are an ancient and diverse class of non-coding RNAs that regulate the expression of downstream genes in response to small molecule ligands (1). Riboswitches typically comprise an upstream highly conserved ligand-binding aptamer domain (AD) coupled to an expression platform (EP) that senses AD binding state and executes a gene-regulatory decision in response (1). Over the past 20 years, considerable progress has been made in understanding AD-ligand interactions, with high-resolution structures of riboswitch ADs revealing diverse and intricate ligand-binding architectures (2–4). In addition, bioinformatic analyses have been used to accurately predict the folding pathways and final structures of ADs using sequence conservation and covariation analysis (5, 6). However, the same bioinformatic analysis is more difficult for EPs because of poor sequence conservation; across evolutionary distances, highly conserved ADs are often coupled with completely different EPs to achieve different modes of gene regulation (7). In general, the mechanisms by which EPs sense AD-ligand binding state and convert that information into a gene expression decision through EP structural switching are poorly understood.

Ligand sensing and switching pose particularly difficult challenges for intrinsic transcriptional riboswitches, which execute their genetic decision during active transcription. The constraint that their genetic decision must be made in a short time window necessitates that they bind ligand cotranscriptionally, which in turn means that the folding of their EP into either an intrinsic terminator or anti-terminator structure must occur within the ms-s timescale of transcription (8). Early work demonstrated that transcriptional riboswitch function is highly dependent upon the rate of transcription (NTP concentration) and transcriptional dynamics (RNAP pausing) (9), but mechanistic details of how EPs fold during the kinetic regime of transcription have been harder to decipher. Recent advances applying diverse biophysical techniques and RNA structure chemical probing methods are beginning to uncover some of these cotranscriptional EP folding mechanisms (10–16). Collectively, these studies are starting to show that many transcriptional ON-riboswitches perform rapid decision-making between two mutually exclusive EP structural states via internal strand displacement mechanisms.

Strand displacement is a conformational switching process in which a substrate duplex of nucleic acids is disrupted by a third invading strand, resulting in a new duplex that contains the invader and one of the original substrate strands (17). This process has been long-appreciated as a mechanism that allows RNA structures with similar free energies to quickly interconvert between each other (18). Recent work is showing that strand displacement is a critical component of transcriptional riboswitch mechanisms. For example, in the fluoride riboswitch a strand displacement process is required to form a terminator hairpin, while fluoride binding stabilizes tertiary interactions that block strand displacement, turning gene expression ON (16). In the case of the purine riboswitch *yxjA*, ligand binding blocks the strand displacement-mediated formation of an anti-terminator hairpin, turning gene expression OFF (11). In both of these riboswitches, a key strand displacement reaction both senses AD-ligand binding state and in the absence of ligand initiates the structural rearrangements necessary to make the corresponding gene expression decision.

Strand displacement has also been widely studied within the field of nucleic acid nanotechnology, where detailed measurements of strand displacement kinetics have been used to uncover sequence and structural features that promote and inhibit the process. For example, mismatches between the invading and substrate strands have been shown to modulate strand displacement reaction kinetics over three orders of magnitude (19–23). The ability to precisely tune strand displacement kinetics has in turn been used to manipulate the function of synthetic nucleic acid systems that utilize strand displacement. For example, mismatches have been applied to substantially decrease leak in catalytic hairpin assembly circuits (24), to bias against undesired strand displacement reactions in complex temporal logic circuits (25), and to enable high fidelity detection of mutations in single-nucleotide-specific programmable riboregulators (26).

The precision with which strand displacement systems can be manipulated, and the fact that a number of transcriptional riboswitches have been shown to utilize strand displacement, together motivate the intriguing hypothesis that riboswitch function can be tuned through EP sequence changes that alter strand displacement kinetics. Here we investigate this hypothesis through systematic studies of the ZTP riboswitch, a model transcriptional ON-switch that has been shown to use strand displacement for its switching mechanism. The ZTP riboswitch senses the metabolic intermediate 5-aminoimidazole–4– carboxamide ribonucleotide (Z) and phosphorylated analogs as indicators of folate stress (27). The highly conserved AD of this riboswitch features a 5’ helix-junction-helix motif (P1-J1/2-P2) and a short 3’ stemloop (P3) connected by a variable linker sequence and an H-type pseudoknot (PK) (Fig. 1A) (4, 28, 29). During transcription in the absence of ligand, a nascent 3’ invading strand disrupts the apo-AD, resulting in the formation of a strong intrinsic terminator hairpin; however, in the presence of Z, ligand-dependent stabilizations of the AD prevent disruption by the invading strand, forcing the EP to follow an alternative folding pathway and allowing RNAP to escape the termination site (Fig. 1A) (15, 27). Thus, the EP both senses AD binding state and commits to a gene expression outcome based on a single attempted conformational switch: disruption of the AD by the invading strand (Fig. 1A) (13, 15, 30).

**Figure 1.**
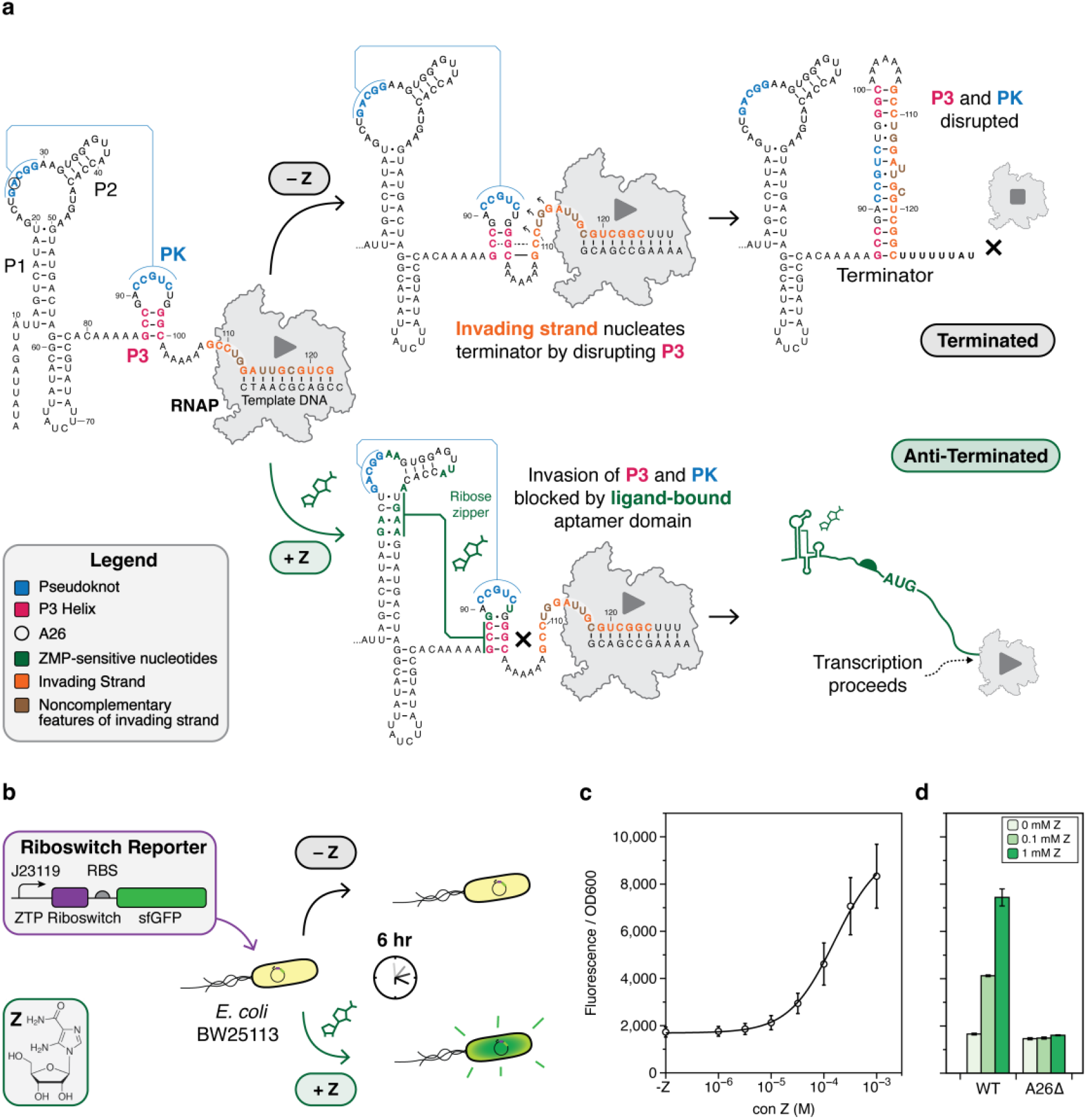
Development of a cellular assay for riboswitch functional mutagenesis. (A) Folding pathway of the *Cbe pfl* ZTP riboswitch highlighting key ligand-dependent structural transitions. In the absence of ligand, the aptamer P3 stem (red) and pseudoknot (PK, blue) are disrupted by the invading strand (orange) of the expression platform via a strand displacement mechanism, forming a terminator hairpin that attenuates transcription. In the presence of Z (green), the aptamer is stabilized by a ligand-dependent hydrogen bonding network that blocks strand displacement, leading to the formation of a non-attenuating alternative structure by the invading strand. The folding pathway was modeled from cotranscriptional SHAPE experiments (15), with nucleotides shown to have altered SHAPE reactivity in the presence of ZMP colored in green. (B) A schematic of the cellular assay used in this work and modified from a similar approach by Kim et al. (27). (C) Dose-response of the wildtype *Cbe pfl* ZTP riboswitch expression construct in *E. coli* with the indicated concentrations of Z supplied to the growth media. (D) A limited dose response of the WT construct and the A26D mutation. Deletion of A26 results in a broken OFF phenotype that no longer responds to Z. Points (C) and bars (D) represent averages from three experimental replicates, each performed with triplicate biological replicates for a total of nine data points (n=9), with error bars representing standard deviation.

Previous work with the *Clostridium beijerinckii* (*Cbe*) *pfl* ZTP riboswitch showed that this disruption of the AD in the absence of Z proceeds via a strand displacement mechanism (15), which has been supported by later smFRET and NMR studies of the *Fusobacterium ulcerans* and *Thermosinus carboxydivorans* ZTP riboswitches, respectively (13, 30). Building off of this work, we used a functional mutagenesis approach with *E. coli in vivo* reporter assays to show that kinetic barriers to strand displacement can be used to tune ZTP riboswitch dynamic range. We first develop an *E. coli in vivo* reporter assay to measure riboswitch function, and subsequently identify sequence and structure features that modulate riboswitch function by tuning the favorability of strand displacement. We next investigated naturally occurring ZTP riboswitch EPs from different organisms. Testing these EP variants in our expression context showed that this strategy may be employed across *Clostridium* to achieve a variety of ON states in different regulatory contexts. As a final step, we synthetically flipped the logic of the *Cbe pfl* ZTP riboswitch by adding a competing strand displacement process, and show that the same kinetic barriers to strand displacement improve dynamic range in this novel context. Together, our findings further elucidate how strand displacement can be manipulated to alter riboswitch dynamic range, suggesting that this could be a mechanism by which evolution tunes riboswitch decision-landscapes, and providing an approach to optimize synthetic riboswitches for biotechnology applications.

## MATERIAL AND METHODS

### Cloning and Plasmid Construction

Each riboswitch reporter plasmid was constructed by inserting the *Clostridium beijerinckii pfl* ZTP riboswitch downstream of the J23119 *Escherichia coli* (*E. coli*) σ^70^ consensus promoter and upstream of a ribosome binding site and the coding sequence for superfolder green fluorescent protein (sfGFP) in a p15A plasmid backbone with chloramphenicol resistance. Riboswitch mutants were generated by inverse polymerase chain reaction (iPCR) using primers ordered from Integrated DNA Technologies (IDT). Using New England Biosciences (NEB) Phusion® High-Fidelity PCR Kit, 50μL PCRs were assembled with 1-10 ng/μL DNA template, 200 μM dNTPs (NEB), 1X Phusion® Buffer, 400 nM each of forward and reverse primers, and 0.25 μL Phusion® polymerase (2,000 U/mL). iPCR products were run on 1% agarose gels to confirm amplification of the desired length products. 0.25 μL DpnI (NEB) was used to digest background plasmid, and PCR cleanup was performed with buffers from the QIAquick PCR Purification Kit (Qiagen) using EconoSpin® All-in-One mini spin columns (Epoch Life Science). Eluted DNA was used to assemble phosphorylation-ligation reactions with 1X T4 DNA Ligase Buffer (NEB), 0.25 μL T4 PNK (NEB), and 0.25 μL T4 DNA ligase (NEB). Ligated products were used to transform NEBTurbo® chemically competent cells, which were recovered in a benchtop shaker at 1000 rpm and 37° C for 1 hour and plated on LB agar plates containing 34 μg/mL chloramphenicol and incubated for 37° C overnight.

The next day, single colonies were picked to inoculate 5 mL LB cultures, which were miniprepped and confirmed by Sanger sequencing (Quintara Biosciences). Correct constructs were re-transformed into 10-beta chemically competent cells (NEB), plated on LB agar plates containing chloramphenicol, and incubated for 37° C overnight. The next day, single colonies were picked to inoculate 6mL LB cultures, which were used to generate duplicate 1 mL glycerol stocks (25% glycerol, stored -80° C) and miniprepped. Sequences were again confirmed by Sanger sequencing (Quintara Biosciences). Sequences of all riboswitch variants can be found in Supplementary Data File 1 along with Addgene accession numbers for select variants.

### In vivo bulk fluorescence reporter assay

Fluorescence assays were performed in *E. coli* BW25113 (Keio collection parent strain). Riboswitch expression plasmids tested in each experiment were transformed into chemically competent *E. coli* BW25113 cells, recovered in a benchtop shaker at 1000 rpm and 37° C for 1 hour, plated on LB agar plates containing 34 mg/mL chloramphenicol, and incubated overnight at 37° C. The following day, plates with *E. coli* colonies were removed from 37° C and stored at room temperature for approximately 7 hours. For each construct, three colonies corresponding to three biological replicates were used to inoculate separate 300 μL cultures of LB media containing 34 mg/mL chloramphenicol in 2 mL 96-well culture blocks covered by breathable seals, which were incubated in a benchtop shaker at 1000 rpm and 37° C overnight. After overnight incubation for 16 hours, cultures from each biological replicate were used to inoculate a subculture. Each subculture was assembled by combining 4 μL of overnight culture with 194 μL freshly prepared M9 enriched media (1X M9 salts, 1 mM thiamine hydrochloride, 0.4% glycerol, 0.2% casamino acids, 2 mM MgSO_4_, 0.1 mM CaCl_2_, 34 mg/mL chloramphenicol) and 2 μL DMSO containing 100x desired final concentration of 5-aminoimidazole-4-carboxyamide-ribonucleoside (Z) (Millipore Sigma) in 2 mL 96-well culture blocks. Separately prepared stocks of M9 salts, glycerol, casamino acids, MgSO_4_, and CaCl_2_ were used for each experimental replicate. After assembly, biological replicate subcultures were covered with breathable seals and incubated in a benchtop shaker at 37° C and 1000 rpm for 6 hours unless otherwise indicated. Samples for fluorescence analysis were prepared after incubation by combining 50 μL of exponential phase cells with 50 μL 1X phosphate buffered saline (PBS) in flat-bottom optically clear plates. Fluorescence measurements (485 nm excitation, 528 nm emission) and OD600 were taken on a Biotek Synergy H1 plate reader with the gain set to 70. All *in vivo* reporter assays were performed with three experimental replicates, each performed with triplicate biological replicates for a total of nine data points (n=9) per mutant.

Data analysis was performed by subtracting the average blank (50 μL M9 enriched media + 50 μL 1X PBS alone) OD600 and fluorescence measurements from the values collected for each sample-containing well. Corrected fluorescence values (arbitrary units) for each well were divided by corrected OD600 values for that same well, to normalize fluorescence values to the number of cells in each well. All statistical analysis (mean, standard deviation, t-tests with Bonferroni correction, curve-fitting) was performed on background corrected, normalized fluorescence values. EC50 values were determined by fitting dose response curves to the formula 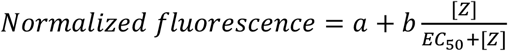 where [Z] denotes ligand concentration, *a* scales the curve to the minimum response and *b* scales the curve to the maximum response. Fitting was performed using DataGraph 5.0.

### Single round in vitro transcription functional assay

Linear templates for *in vitro* transcription (IVT) were generated from plasmids via PCR using primers ordered from Integrated DNA Technologies (IDT). Using New England Biosciences (NEB) Phusion® High-Fidelity PCR Kit, 2 mL PCRs were assembled with 1-10 ng/μL DNA template, 200 μM dNTPs (NEB), 1X Phusion® Buffer, 400 nM each of forward and reverse primers, and 5 μL Phusion® polymerase (2,000 U/mL). PCR products were run on 1% agarose gels to confirm amplification of the desired length product, and PCR cleanup was performed with buffers from the QIAquick PCR Purification Kit (Qiagen) using EconoSpin® All-in-One mini spin columns (Epoch Life Science). IVT linear template was eluted from spin columns using 50 μL UltraPure™ DNase/RNase-Free Distilled Water (Invitrogen) and quantified using a NanoDrop™ One Spectrophotometer (Thermo Scientific).

25 μL reactions were prepared containing 90 nM linear DNA template, 0.5 μL 50x Z in DMSO, transcription buffer (20 mM Tris-HCl, pH 8.0, 0.1 mM EDTA, 1 mM DTT, 5 mM MgCl_2_ and 50 mM KCl), 0.1 mg/ml BSA (NEB), and 1.25 μL of *E. coli* RNAP holoenzyme (NEB). The reaction tube was incubated at 37° C for 10 minutes to form open transcription complexes, and then started with NTPs (final concentration of 400 μM ATP, GTP, CTP, UTP) and rifampicin at 10 μg/mL. Transcription was allowed to proceed for 30 s at 37° C and then stopped by the addition of 75 μL TRIzol™ Reagent (Invitrogen).

RNA extraction and purification was performed by adding 20 μL chloroform (Millipore Sigma) and spinning for 5 min at 4° C. 70 μL of the aqueous phase was transferred to a new tube containing 50 μL isopropanol (Millipore Sigma) and 1 μL glycogen coprecipitant (Invitrogen). Samples were incubated at room temperature for 20 min, pelleted by spinning at 4° for 45 min, aspirated, and washed with 500 μL 70% ethanol. Dried pellets were resuspended in 1X TURBO™ DNase Reaction Buffer (Invitrogen) and 1 μL TURBO™ DNase and incubated at 37° for 1 hr. Following digestion, RNA extraction and purification was performed using 150 μL TRIzol™ Reagent (Invitrogen) and 40 μL chloroform (Millipore Sigma). 140 μL of the aqueous phase was extracted and transferred to a new tube containing 100 μL isopropanol and 1 μL glycogen coprecipitant (Invitrogen). RNA was pelleted by spinning at 4° C for 45 min, and aspirated and washed with 70% ethanol. Dried pellets were resuspended in 10 μL UltraPure™ DNase/RNase-Free Distilled Water (Invitrogen) and 10 μL 2X RNA Loading Dye (NEB), denatured at 95° for 5 min, and run on an 8% denaturing gel made using reagents from the SequaGel^®^ UreaGel™ System (National Diagnostics). Gels were stained with SYBR™ Gold Nucleic Acid Gel Stain (Invitrogen) and imaged on a ChemiDoc™ Touch Imaging System (BioRad). Relative band intensity was determined using Image Lab 6.1 (BioRad).

## RESULTS

### Developing an *in vivo* reporter assay to characterize riboswitch function

We began by developing an expression construct that would allow us to characterize riboswitch function *in vivo* using cellular gene expression assays. The expression construct consisted of the well-characterized *Cbe pfl* ZTP riboswitch sequence placed downstream of a J23119 constitutive *E. coli* σ70 promoter as previously described (15), followed by a strong ribosome binding site and a superfolder GFP (sfGFP) coding sequence (Fig. 1B, Supplementary Data File 1). This construct was then placed within a p15A plasmid backbone with chloramphenicol resistance. Following an experimental design by Kim and colleagues (27), who used direct treatment of *E. coli* with Z to induce a β-galactosidase expression cassette, we transformed this plasmid into *E. coli* strain BW25113, picked colonies, grew overnight cultures, and sub-cultured with and without Z added directly to the media (Methods, Fig. 1B). Endpoint bulk fluorescence and optical density (OD600) measurements were made after 6 hours of subculture in the presence or absence of different amounts of Z. We selected 6 hours as our endpoint measurement because this time point showed robust fold-changes between the no ligand and saturating ligand (1 mM) conditions (Fig. S1A). Under these conditions, we found that the addition of Z did not affect growth rate (Fig. S1B), and that this riboswitch expression construct responded comparably to Z and ZMP (Fig. S1C). Using this *in vivo* reporter assay, we characterized riboswitch dose-response by adding a range of Z concentrations to the media (Fig. 1C), which showed similar dose-response characteristics to those observed using *in vitro* transcription (15).

Using the *in vivo* assay, we next characterized a riboswitch mutant broken in the OFF functional state. Following previous mutational analysis of the PK region which rendered the riboswitch unresponsive to induction by ZMP (15), we selected A26Δ as the OFF control. Characterization of this mutant with several Z concentrations showed the expected effects (Fig. 1D). All subsequent mutational analysis was also carried out in the A26Δ background as a negative control, which allowed us to distinguish between mutations that truly affect the anti-termination decision and mutants that affect baseline termination efficiency alone. Taken together, these results establish an *in vivo* assay for functional characterization of *Cbe pfl* riboswitch mutants.

### *Cbe pfl* riboswitch EP sequence and structure features tune dynamic range

A key goal of this study was to interrogate how specific features of the *Cbe pfl* ZTP EP determine riboswitch function *in vivo*. The key conformational change enabling switching behavior in the cotranscriptional folding pathway of the ZTP riboswitch is strand displacement of the AD P3 helix by the 3’ invading strand (Fig. 2A). This step is necessary in order to disrupt the AD and form the terminator in the OFF state, but must be blocked in order to allow transcription to proceed in the ON state. How this strand displacement process is governed by EP sequence and structure remains poorly understood. We therefore sought to interrogate this critical conformational change by identifying nucleotides in the terminator invading strand that are important for regulatory function. Previous smFRET experiments with the *Fusobacterium ulcerans* ZTP riboswitch showed that mutating the first three nucleotides of the invading strand to A severely delayed formation of the terminator hairpin (13), inspiring us to interrogate the invading strand with fine-grained functional mutagenesis.

**Figure 2.**
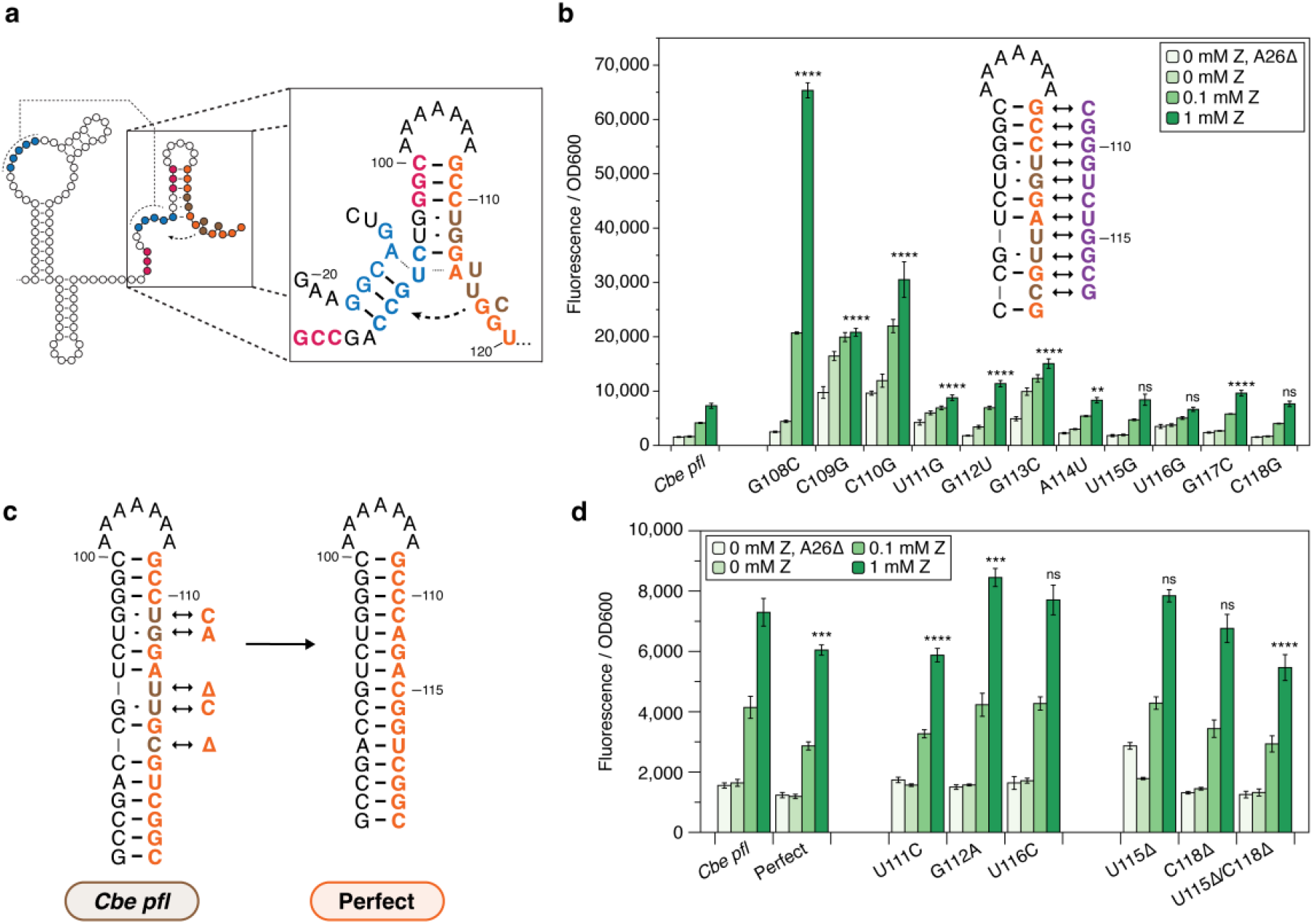
Noncomplementary elements in the expression platform tune the ON state of the *Cbe pfl* ZTP riboswitch. (A) Diagram of the strand displacement process during terminator formation. The invading strand (orange) nucleates the terminator hairpin by base pairing with the 3’ nucleotides of the P3 stem (red), enabling branch migration through the pseudoknot (PK, blue). Noncomplementary elements (mismatches, bulges) of the invading strand are shown in brown. (B) Mismatch scan of the invading strand showing position-dependent effects of mismatch mutations on fluorescence, mostly affecting expression when ZTP is present. Asterisks indicate the significance threshold α for pairwise t-tests between the observed fluorescence from a particular mutation vs. that from the wildtype construct at the 1mM ZTP condition. (C) Diagram highlighting the noncomplementary elements of the natural invading strand (brown), and the mutations performed to generate a perfectly complementary invading strand. (D) Mutational scan to eliminate noncomplementary elements individually, or all at once (perfect). Noncomplementary elements are grouped into wobble pairs (center) or bulges (right). Asterisks indicate the significance threshold α for pairwise t-tests between the observed fluorescence from a particular mutation vs. that from the wildtype construct at the 1mM ZTP condition. Bars in (B) and (D) represent averages from three experimental replicates, each performed with triplicate biological replicates for a total of nine data points (n=9), with error bars representing standard deviation. T-test significance thresholds are for pairwise (two-tail, heteroscedastic) tests with values prior to Bonferroni correction of: * = 0.05, ** = 0.01, *** = 0.001, **** = 0.0001, ns = not significant.

We disrupted the formation of each base pair of the terminator by mutating each nucleotide of the invading strand in two ways: either with a mismatch mutation (Fig. 2B), or by deleting each nucleotide (Fig. S2A). Gene expression analysis of these variants in the presence of varying amounts of Z showed that the base pair interactions closest to the strand displacement nucleation site (top of the terminator hairpin) have the biggest effect on riboswitch dynamic range, consistent with the observations of Hua et al. (13). Specifically, we found that mutating the terminator nucleating base pair (C100:G108) to either a mismatch (G108C) or a deletion (G108Δ) resulted in a significantly higher dynamic range than the wildtype riboswitch (Fig. 2B, S2A), suggesting that the formation of the wildtype GC pair in this position has a dampening effect that limits the natural ON state. Sequence changes further down the terminator hairpin have progressively smaller effects on dynamic range (Fig. 2B, S2A). Surprisingly, the C118G mismatch mutation, which should disrupt a previously described RNAP pause site, did not result in any significant change in ON-state gene expression (Fig. 2B). Other C118 mutants did not show large differences in sensitivity or dynamic range compared to wild type (Fig. S2B-D), indicating that pause site 2 does not play an important role in riboswitch function. Taken together, these results align with previous observations from DNA nanotechnology that mismatch mutations closer to the strand displacement initiation site have larger effects on the strand displacement rate than more distal mismatch mutations (20–22).

Based on these results, we reasoned that noncomplementary base pairing features present in the natural sequence of the invading strand could pose kinetic barriers to strand displacement, disfavoring the formation of the terminator hairpin. Specifically, the wildtype *Cbe pfl* invading strand features five non-complementary elements relative to the sequence that it invades: three wobble pairs and two single-nucleotide bulges (Fig. 1A, 2C; brown). To test if these wobble pairs and bulges moderate the kinetics of strand displacement to achieve higher baseline expression and/or higher ON state, we ‘corrected’ the invading strand to perfectly complement the substrate strand contained in the AD (Fig. 2C). We also characterized the effect of making each of these corrective mutations individually. We observed that the perfectly complementary invading strand resulted in significantly lower ON state than wildtype *Cbe pfl* (Fig. 2D), and of the five individual corrective mutations, the correction of the wobble pair most proximal to the strand displacement nucleation site (U111C), decreased the ON state the most (Fig. 2D). While deleting either the U115 or C118G bulges individually did not significantly affect the ON state, deleting both bulges simultaneously resulted in a substantial decrease in baseline expression and ON state compared to the wild type (Fig. 2D). Interestingly, the G112A corrective mutation increased ON-state fluorescence compared to the wild type, suggesting that the natural U96:G112 wobble pair facilitates more efficient strand displacement through P3 than a Watson-Crick pair at this position. In total, these results show that the *Cbe pfl* ZTP riboswitch EP contains sequence features that tune dynamic range by moderating the kinetics of strand displacement.

### *Clostridium* genus ZTP riboswitches contain EP elements that tune dynamic range

Having established that the *Cbe pfl* ZTP riboswitch uses noncomplementary elements to tune ON-state gene expression by moderating the kinetics of strand displacement, we hypothesized that other *Clostridium* genus ZTP riboswitches may employ similar mechanisms to match riboswitch function to cellular needs. To investigate this hypothesis, we first performed a bioinformatic analysis of EP sequence variation within ZTP riboswitches. All *Clostridium* genus ZTP riboswitch sequences from the RFam database (73 sequences) were aligned, and first narrowed to include only those sequences that contain a putative terminator poly-U tract of at least 5 Us and an identical P3 sequence to the *Cbe pfl* riboswitch, which resulted in 18 curated sequences. We next identified those sequences which feature a loop + invading strand architecture similar to the *Cbe pfl* riboswitch based on secondary structure prediction of the terminator hairpin using NUPACK (15 sequences) (31).

From this set, we selected a subset of invading strand variants that i) possessed a unique invading strand, and ii) which natively regulate a gene also regulated by at least one other riboswitch in the subset, for a final subset of 12 *Clostridium* genus ZTP riboswitches, including a second ZTP riboswitch from *C. beijerinckii*. For each subset member, we created a chimeric riboswitch consisting of the *Cbe pfl* AD (including the P3 sequence shared among all sequences) and the invading strand of the variant of interest (Fig. 3A). Cellular gene expression assays of the 8 functional chimeras using a range of ZTP concentrations showed that variations in the invading strand sequence (Supp Data Table 1) led to differences in functional properties of the chimeric riboswitches (Fig. 3A).

**Figure 3.**
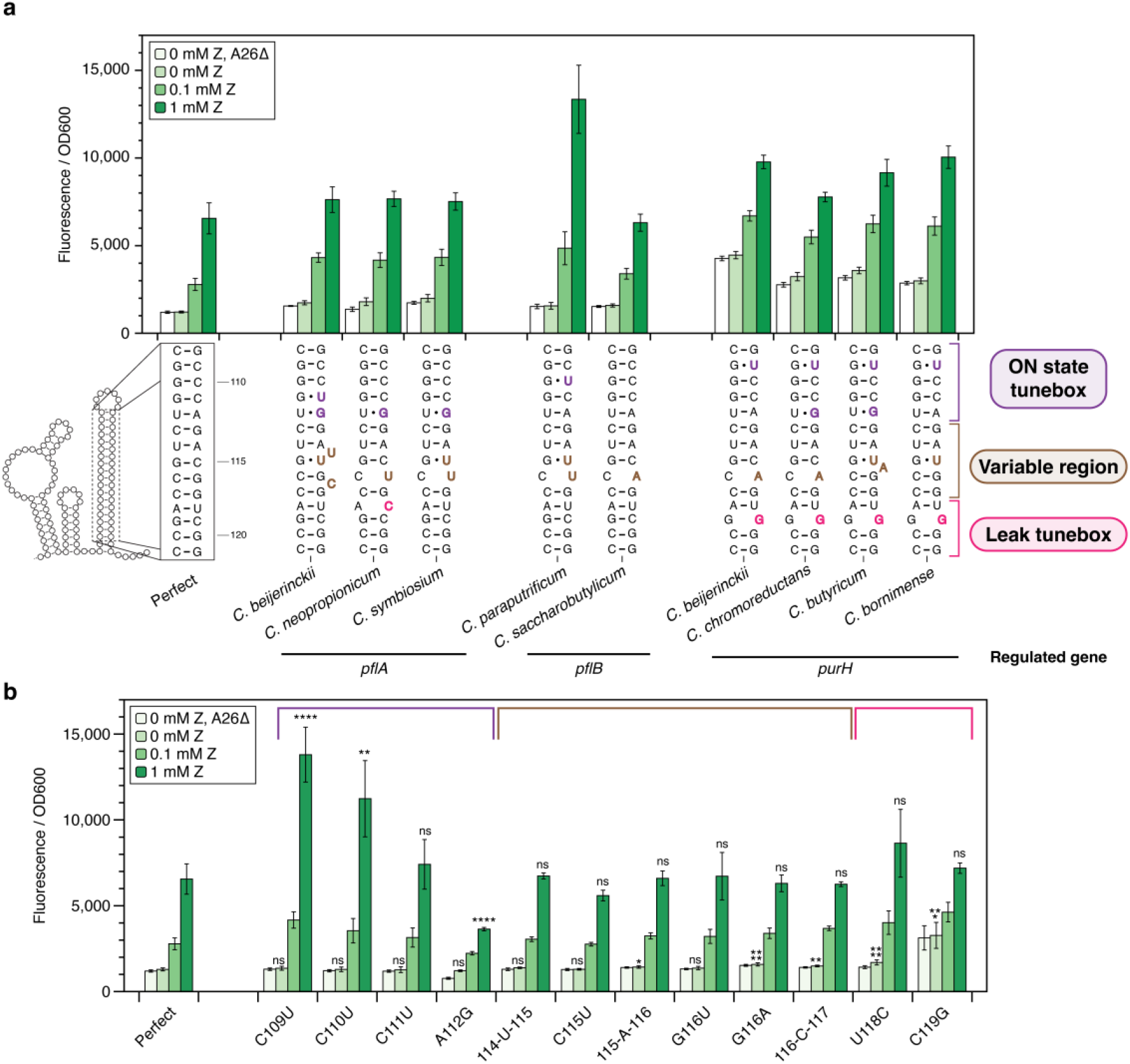
Other *Clostridium* ZTP riboswitch expression platforms tune ON and OFF states with kinetic barriers to strand displacement. (A) Gene expression characterization of chimeric riboswitches, constructed by fusing the *Cbe pfl* riboswitch aptamer sequence (containing the 5’ portion of the expression platform) with the 3’ invading strand of the expression platform of different *Clostridium* species. Putative terminator hairpin secondary structures (noncomplementary elements in bold) are below observed gene expression under varying ZTP concentration conditions. Sequences are grouped by the gene they regulate in their native genomic context (Supplementary Table 1). Bolded sequence elements are colored according to their position within the annotated regions of the invading strand. (B) Functional characterization of all individual noncomplementary elements from natural *Clostridium* ZTP riboswitch expression platforms characterized in (A) in the context of the perfectly complementary terminator (Fig. 2C). Insertion mutations are bracketed by position numbers. Based on the magnitude of functional effects, these mutations are divided into three functional domains: the ON-state tunebox (purple), the variable region (brown), and the leak tunebox (fuchsia). Bars represent averages from three experimental replicates, each performed with triplicate biological replicates for a total of nine data points (n=9), with error bars representing standard deviation. Asterisks indicate significance threshold α for pairwise t-tests (two tailed, heteroscedastic) between the observed fluorescence from a particular mutation vs. that from the *pfl* construct with perfect expression platform pairing at the ZTP condition indicated on the bar (prior to Bonferroni correction: * = 0.05, ** = 0.01, *** = 0.001, **** = 0.0001.

Consistent with our previous observations, invading strands with kinetic barriers to strand displacement near the top of the terminator hairpin showed higher ON-state fluorescence than the perfectly complementary invader (Fig. 3A). To validate these results, we determined the functional effect of all non-complementary sequence elements from these riboswitches in the perfect invading strand background (Fig. 3B). These results allowed us to demarcate regions of the invading strand responsible for tuning the ON state and the leak level, as well as a variable region relatively insensitive to mutation (Fig. 3A, B).

Interestingly, we observed that invading strands from riboswitches that regulate the same genes result in functionally similar chimeric riboswitches *in vivo* (Fig. 3A). For example, *purH*-regulating riboswitches all show elevated ON-state fluorescence relative to the perfectly complementary invader, which can be attributed to the common C110U wobble pair mutation, and all these chimeras also show elevated baseline expression compared to the perfectly complementary invader (Fig. 3A). This increase in leak is attributable the presence of a G:G mismatch near the base of the terminator hairpin (Fig. 3B), consistent with known structure rules dictating intrinsic terminator strength (32, 33). In the case of ZTP riboswitches that regulate *pflA*, the resulting chimeric riboswitches were nearly functionally identical, despite deviations in the sequence of the invading strands (Fig. 3A). Taken together, these findings suggest that natural ZTP riboswitches could have evolved to exploit strand displacement kinetics to tune expression in different regulatory contexts.

### Kinetic barriers to strand displacement tune ON-state fluorescence

Taking inspiration from the natural strategy employed at least by *Clostridium* ZTP riboswitches to tune ON-state gene expression with noncomplementary features in the invading strand, we next sought to perform fine-grained analysis of how different mutation types at different positions in the invading strand alter dynamic range (Fig. 4A). We chose to perform this mutagenesis analysis in the context of the perfectly complementary invader (Fig. 2C) so we could measure the effect of each mutation in isolation.

**Figure 4.**
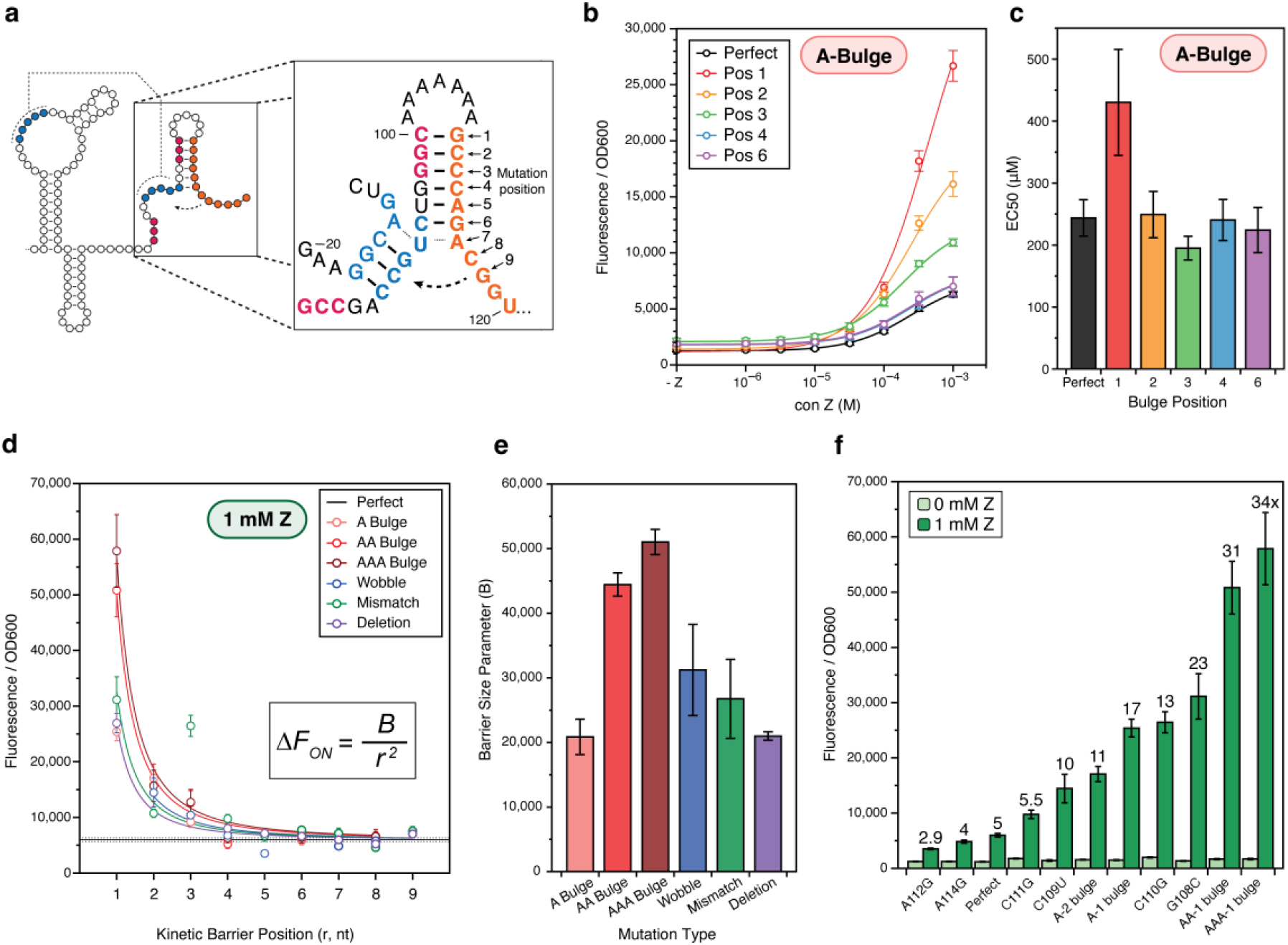
Kinetic barriers to strand displacement enable fine-grained control of riboswitch dynamic range. (A) Diagram of the strand displacement process during terminator formation in the context of the perfectly paired terminator (Fig. 2C). The invading strand (orange) nucleates the terminator hairpin by base pairing with the 3’ nucleotides of the P3 stem (red), enabling branch migration through the PK (blue). Positions mutated in subsequent panels are indicated by a numerical index. Bulge mutations were inserted immediately after the nucleotide identified by the position index (e.g. A bulge at position 3 denotes an insertion between C110 and C111) (B) Dose response curves for various A-bulge mutants. (C) EC50s extracted from sigmoidal fits of the data in panel B showing the effect of A-bulge mutations on riboswitch sensitivity. Error bars indicate standard error of the fit parameter. (D) The measured ON state of gene expression assays with 1 mM ZTP from scans with different mutations including bulge insertions (A, AA, AAA), wobble pairs, mismatch pairs and deletions, each at the different indicated positions within the terminator. The perfectly complementary expression platform fluorescence is shown as a dashed line. Full data in Fig. S3 A-E. Data are fit to the inset equation, where Δ*F* is the increase in ON-state fluorescence for each mutant compared to the perfect terminator ON-state fluorescence, and the parameter B characterizes the kinetic ‘barrier size for strand invasion. (E) The kinetic barrier size parameter, B, extracted from the fits plotted in (D), showing the relative effects of different mutations types on ON-state fluorescence. Error bars indicate standard error of the fit parameter. (F) Gene expression characterization of select kinetic barrier mutants showing widely diverging dynamic range. Fold-change between 1mM and 0mM ZTP conditions is indicated by a label above each mutant. Error bars indicate standard deviation. Points in (B) and (D) and bars in (F) represent averages from three experimental replicates, each performed with triplicate biological replicates for a total of nine data points (n=9), with error bars representing standard deviation.

We first tested the addition of single-nucleotide bulges at different possible positions withing the invading strand in order to slow strand displacement and thus disfavor terminator formation. As expected, we found that these mutations resulted in an increase in ON-state gene expression in a position-dependent manner (Fig. 4B). Interestingly, the A-bulge at position 1 affected EC50, indicating that early events in the strand displacement pathway could affect sensitivity; though this effect was not observed for any other position (Fig. 4C).

To extend this analysis, we performed similar mutational scans by inserting di- and tri-nucleotide bulges, which we expected to pose larger barriers to strand displacement than a single-nucleotide bulge, as well as mismatches, wobble pairs, and deletions. All of these mutational scans resulted in similar position-dependent increases in ON-state fluorescence relative to the perfectly complementary invader scaffold, though they appeared to have a different magnitude of effect depending on the type of mutation introduced (Fig. 4D, S3A-E).

To better understand the difference in magnitude between different types of mutations at a phenomenological level, we fit the position-dependent ON state values to the power law curve 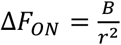, where Δ*F*_*ON*_ is the increase in ON-state fluorescence for each mutant compared to the perfect terminator ON-state fluorescence, *r* is the position of the mutation in nucleotides from the strand displacement nucleation site, and *B* describes the magnitude of the decay trend (Fig. 4D). Plotting the fitted *B* values for each type of mutation showed that the trend was single nt bulge = deletion < mismatch ≤ wobble < double nt bulge < triple nt bulge (Fig. 4E). This aligns with the interpretation that the larger the barrier to strand displacement that each mutation produces, the larger the phenomenological *B* value.

Overall, these results show that introducing kinetic barriers to strand displacement by mutating the invading strand enables fine-tuning of riboswitch dynamic range *in vivo* from 2.4 to 34 fold (Fig. 4F), and larger kinetic barrier mutations placed closer to the initiation of strand displacement cause the largest effect.

### Kinetic strand displacement barriers can be used to tune the dynamic range of a synthetically flipped-logic ZTP riboswitch

As a final demonstration of the importance of strand displacement barriers for tuning riboswitch decision-making, we sought to apply our understanding from the studies above to tune the functional performance of a version of the *Cbe pfl* ZTP riboswitch that features an additional competing strand displacement process. In a recent study of the *yxjA* purine transcriptional OFF-riboswitch, it was discovered that this riboswitch flips the typical transcriptional ON regulatory logic through the presence of an additional intermediate structure (11). Inspired by this observation, we inverted the regulatory logic of the *Cbe pfl* ZTP riboswitch by inserting an 8-nt ‘flipping strand’ after the intrinsic terminator hairpin (Fig. 5A). In the absence of ligand, this flipping strand is designed to act as a spacer between the wildtype intrinsic terminator hairpin and the poly-U tract, thus ablating termination and producing an ON regulatory state. In the presence of ligand however, the flipping strand was designed to base pair with the remaining portion of the wildtype invading strand left over from failed strand displacement of the AD, creating a new synthetic intrinsic terminator hairpin adjacent to the poly-U tract and resulting in an OFF regulatory state. As expected, this synthetic OFF-riboswitch functions at the level of transcription, though it exhibits poorer dynamic range than the wildtype ON-switch *in vitro* (Fig. 5B) and *in vivo* (Fig. 5C).

**Figure 5.**
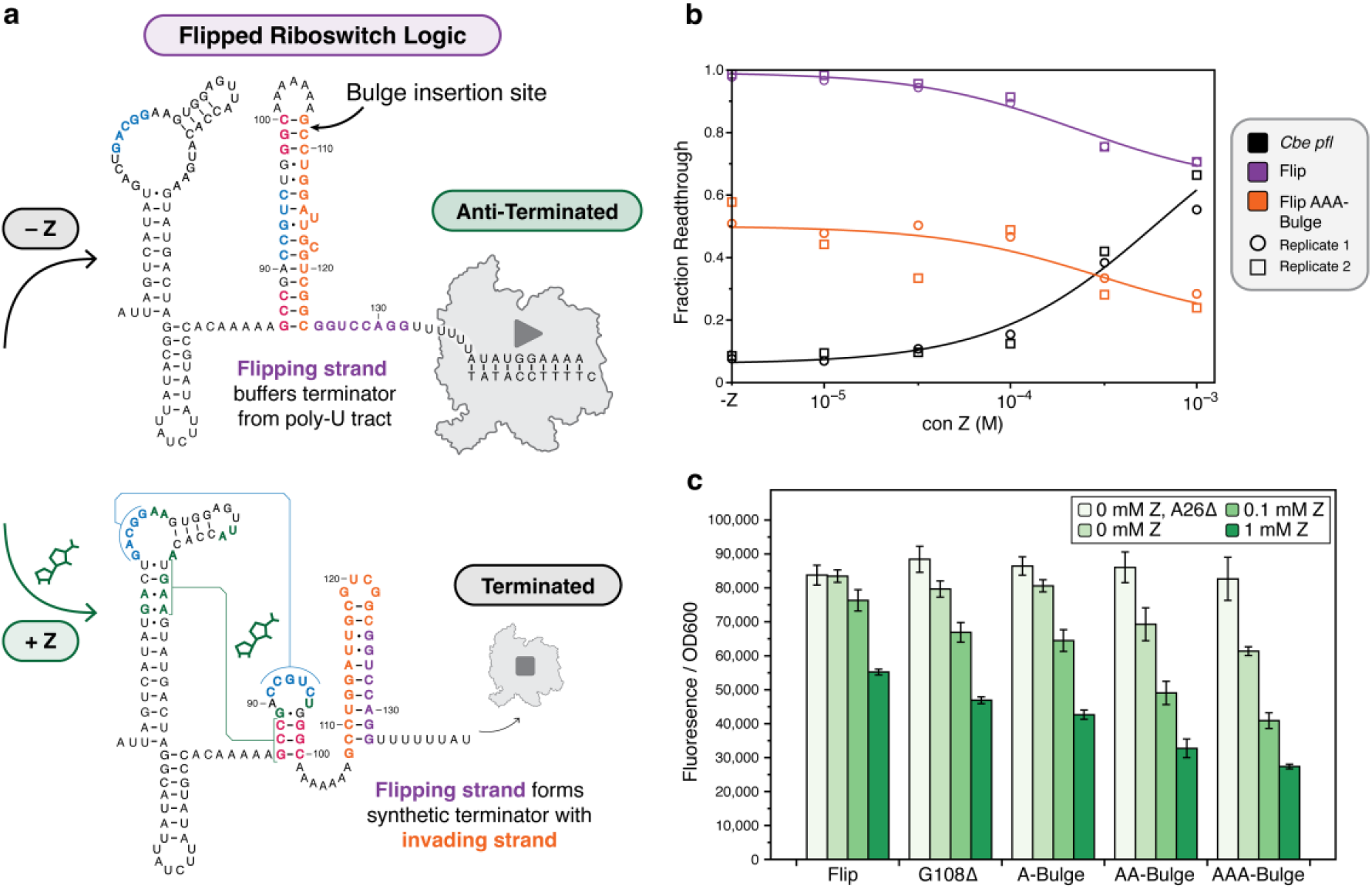
Kinetic barriers to strand displacement tune the dynamic range of a synthetic ‘flipped-logic’ ZTP riboswitch. (A) Schematic detailing the approach for inverting the logic of the *Cbe pfl* ZTP riboswitch from an ON-switch to an OFF-switch. In the absence of ligand, an eight nucleotide insertion, named the “flipping strand” (purple), is designed to ablate intrinsic termination by increasing the spacer between the ZTP riboswitch terminator hairpin and its poly-U tract. In the presence of ligand, however, the flipping strand is designed to form a synthetic terminator with the portion of the expression platform that results from unsuccessful strand displacement of the aptamer. Areas in the expression platform where sequence deletions or bulge insertions can tune dynamic range are indicated. (B) Single round *in vitro* transcription results validating the functionality of the ‘Flip’ design and the functional improvement imparted by a tri-nucleotide bulge between G108 and C109. Source gels are available in Supplementary Figures S4-7. (C) *In vivo* validation of the ‘Flip’ design showing successive improvements in dynamic range when successively larger kinetic barriers to strand displacement of the aptamer are added to the expression platform at the position indicated in (A). Bars in (C) represent averages from three experimental replicates, each performed with triplicate biological replicates for a total of nine data points (n=9), with error bars representing standard deviation.

We reasoned that the observed high leak level in the ligand-induced OFF state was the result of an inherent bias toward the formation of the wildtype terminator hairpin, which in this flipped context would result in a bias towards the ON state. Based on our understanding of strand displacement barriers, we hypothesized that inserting kinetic barriers into the invading strand would reduce this bias and thereby improve dynamic range. In this context, the G108Δ deletion, which increases the ON state of the *Cbe pfl* riboswitch >3x (Fig. S2A), significantly decreases the leak of the flipped-logic OFF-switch (Fig. 5C). Larger kinetic barriers, including a di- or tri-nucleotide bulge in between G108 and C109, result in greater decreases in OFF state leak (Fig. 5C), indicating that kinetic barrier size is correlated with the resulting dynamic range.

These results demonstrate the ability to flip the logic of transcriptional ON-riboswitches into OFF-riboswitches using synthetic sequences, and further show how the application of simple mutations to slow strand displacement at key points of the riboswitch mechanism can be used to enhance the dynamic range off synthetic transcriptional OFF riboswitches as well.

## DISCUSSION

There is a growing appreciation for the apparent ubiquity of DNA and RNA mechanisms that feature strand displacement-mediated conformational switching (34). Processes at the heart of genome maintenance and gene expression such as homologous recombination, spliceosomal activation and large ribosomal subunit maturation are thought to be facilitated by strand displacement-mediated rearrangements (17, 35, 36). However, there are still large gaps in our knowledge of how biology tunes these processes *in vivo* (17, 34). Recent work has shown that several classes of transcriptional riboswitches feature an internal strand displacement process at the core of their folding pathways, which enables EPs to both sense AD-ligand binding state and commit to a gene-regulatory decision based on the outcome of this attempted conformational switch. In this work, we observed that the *Cbe pfl* ZTP riboswitch balances this cotranscriptional conformational switching with a combination of sequence and structure features that both favor and disfavor efficient strand displacement. On the one hand, noncomplementary sequence elements in the EP invading strand act to moderate strand displacement kinetics and increase ON-state gene expression. On the other hand, the wildtype EP forms several strong nucleating base pairs at the initiation of strand displacement, the disruption of which ‘unlocks’ drastic increases in dynamic range. Similar findings were observed in a recent study of the *Fusobacterium ulcerans* ZTP riboswitch, which showed that mutations to the EP invading strand severely slowed the rate of terminator formation *in vitro*, supporting our conclusion that modulation of strand displacement kinetics tunes riboswitch conformational switching (13).

A quantitative look at which types of sequence variations led to the largest changes in riboswitch function shows that there may be limits to how much tuning strand displacement can alter riboswitch function. While most kinetic barrier mutations had minimal impact on riboswitch leak, the kinetic barriers with the largest B values in this study, di- and tri-nucleotide bulges, produced large increases in uninduced gene expression when placed at positions proximal to the strand displacement nucleation site (Fig. S3B). Interestingly, these increases in leak are not observed in the PK-incompetent A26Δ background (Fig. S3A), suggesting that sufficiently large kinetic barriers to EP strand displacement can disfavor formation of the terminator hairpin enough to allow a subpopulation of apo-ADs to withstand displacement long enough to yield an anti-termination decision. Taken together, these results point to inherent limits in the switching efficiency achievable by tuning strand displacement.

This work also sheds light on two important general questions in riboswitch biology: i) Why are EPs so poorly conserved relative to ADs, and ii) How can bacteria regulate multiple genes at different baseline levels with the same riboswitch class? In this work, we observed that EPs from different *Clostridium* species with multiple different point mutations can achieve nearly identical functional profiles (the same baseline and ON-state gene expression). A more detailed analysis shows that the ZTP riboswitch EP contains distinct regulatory regions that affect different aspects of gene expression. Overall, this supports the view that poor EP conservation arises partly because there are many possible EP sequences that can meet evolutionary selection pressures on function. Importantly, the mechanistic basis of how sequence variation in the EP leads to leak and ON-state tuning requires only the *disruption of* base pairing to provide a kinetic barrier to strand displacement, in contrast to mutational covariation needed to preserve RNA structures in ADs during riboswitch evolution. Thus EP sequence variation may offer a simpler evolutionary route to tuning riboswitch function than AD sequence variation. We hope these observations can be used to inform new bioinformatic analysis that can further uncover patterns in EP sequence evolution.

Our work in creating a synthetically flipped OFF ZTP riboswitch also synergizes with recent studies of natural transcriptional OFF-riboswitches. Recently we showed that the *yxjA* purine OFF-switch contains an additional ‘central helix’ intermediate structure, formed cotranscriptionally in between the AD and EP, that serves to introduce an additional competing strand displacement process that inverts the regulatory logic from the typical ON adenine/guanine riboswitch mechanism (11). Work from Schwalbe and colleagues also recently highlighted a contrasting pair of cyclic-di-nucleotide-sensing riboswitches with opposite transcriptional logic, with the Cd1 OFF-switch EP featuring an additional competing structure relative to the simpler pilM ON-switch, resulting in inverted regulatory logic (37). In this context, our introduction of a synthetic ‘flipping’ strand converts the ZTP terminator hairpin into a central-helix-like element, and introduces an additional competing strand displacement process that flips the regulatory logic. This points to a potentially generalizable mechanism of converting transcriptional ON-switches into OFF-switches through the insertion of additional sequences, and vice versa, which could have implications for riboswitch evolution.

Finally, we believe this work has implications for biotechnological applications of riboswitches, and adds to the list of examples of tools that use the EP as a programmable scaffold to modulate riboswitch dynamic range (38). As ADs are extremely sensitive to mutation, and among the most highly conserved sequences known (39), engineering EPs appears to be a more fruitful approach of optimizing natural and synthetic riboswitches for diverse applications in diagnostics, gene-regulatory circuits, or reporters (40–42).

## Supporting information

Supplemental Data File 1 Sequences

Supplemental Data File 2 All Source Data

## SUPPLEMENTARY DATA

Supplementary Data are available at NAR online.

## ACKNOWLEDGEMENTS

We wish to acknowledge the helpful insight of all members of the Lucks lab, especially Dr. Katherine Berman and Dr. Luyi Cheng for discussions about flipping riboswitch logic, Reese Richardson and Laura Hertz for advice in sequence analysis, and Edric Choi for critical review of figures. We also wish to thank Dr. Narasimhan Sudarsan for helpful advice about selecting strains for *in vivo* riboswitch assays.

## FUNDING

This work was supported by National Institutes of Health Research Project grants [R01GM130901] and in part by a National Institutes of Health Training Grant [T32GM008382 to D.Z.B] through the Northwestern University Molecular Biophysics Training Program.

## CONFLICT OF INTEREST

The authors declare that they have no known competing financial interests or personal relationships that could have appeared to influence the work reported in this paper.

## Supplementary Info for

**Supplementary Figure 1.**
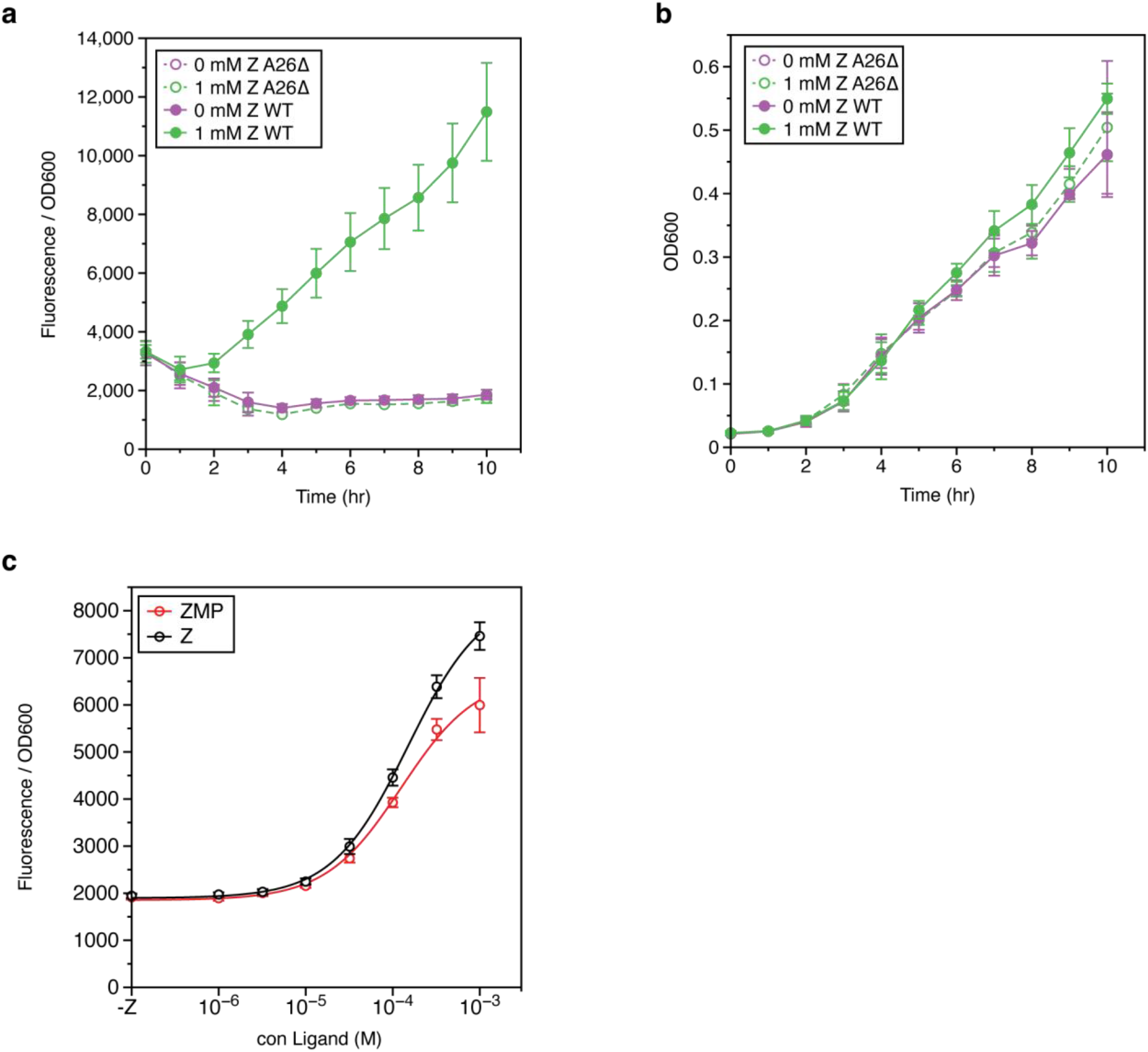
Additional validation for *in vivo* reporter assay. (A) Time-course measurements of GFP fluorescence under the regulation of the *Cbe pfl* ZTP riboswitch show that by 6 hrs robust dynamic range is observed between the induced (1 mM Z) and uninduced (0 mM Z) subcultures. (B) Time-course measurements of subculture optical density shows that Z-induction has little effect on growth rate. (C) Dose-response curve at 6 hrs comparing response when the *Cbe pfl* ZTP reporter cassette is induced with Z or its monophosphorylated analog ZMP. Both curves show similar sensitivity and dynamic range, validating the use of Z in subsequent experiments. Data points represent averages from three experimental replicates, each performed with triplicate biological replicates for a total of nine data points (n=9), with error bars representing standard deviation.

**Supplementary Figure 2.**
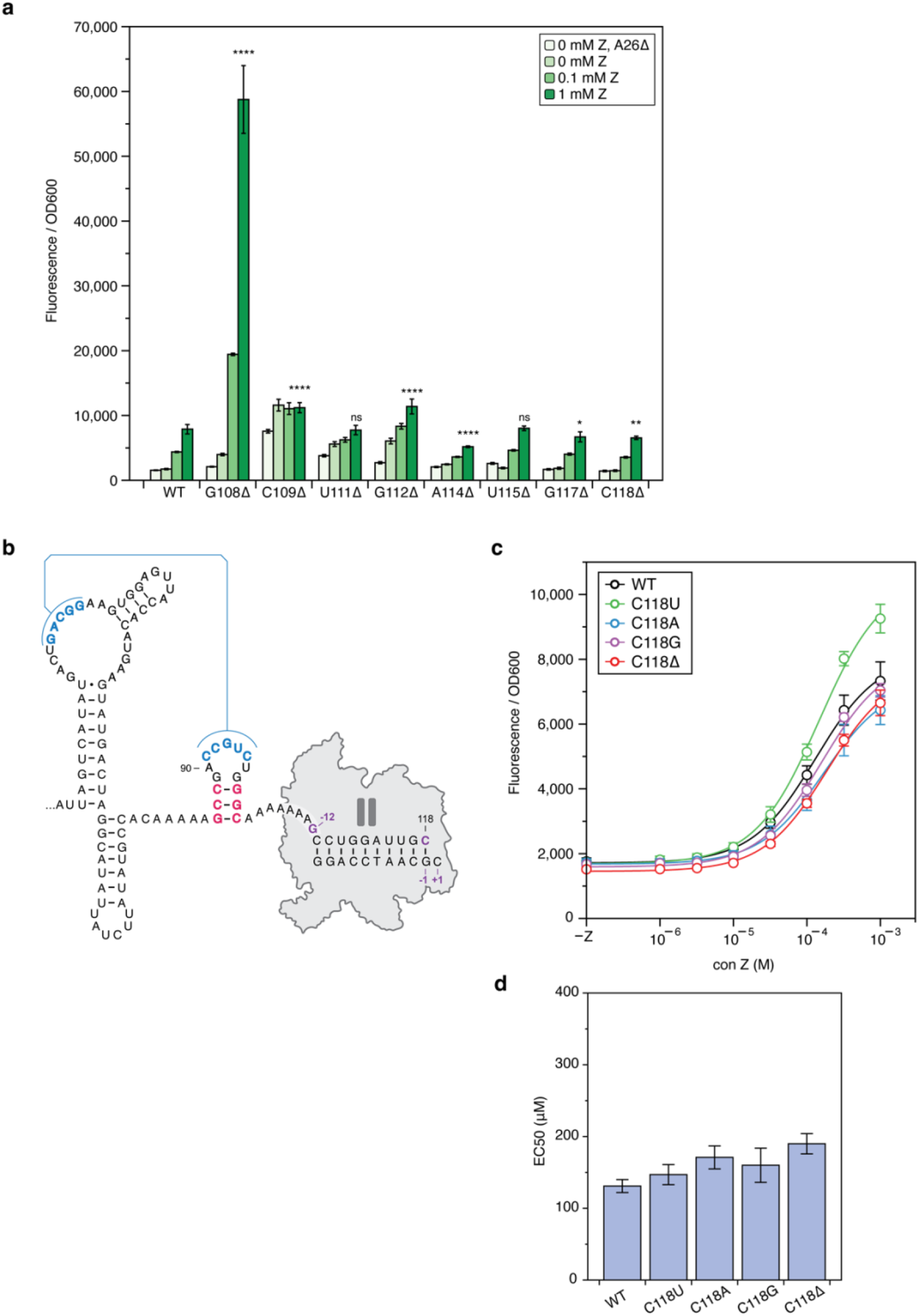
Noncomplementary elements mediate tuning of dynamic range. (A) Deletion scan of the invading strand shows position-dependent effects of deletion mutations on ON-state fluorescence. (B) Structure diagram of the C118 pause site 2. The ligand-binding competent aptamer is able to form. (C) Mutations designed to disrupt the C118 RNAP pause site 2 do not result in large changes to dynamic range. Data was fit according to a sigmoidal shape (see Methods). (D) EC50 values from the fits in (C) showing little change in riboswitch sensitivity from these variants. Bars in (A), and points in (C) represent averages from three experimental replicates, each performed with triplicate biological replicates for a total of nine data points (n=9), with error bars representing standard deviation. Bars in (D) represent values for the parameter EC50 (For equation see Methods), with error bars indicating standard error of the fit parameter. T-test significance thresholds in (A) are for pairwise (two-tail, heteroscedastic) tests between the 1mM and 0mM Z conditions, with values prior to Bonferroni correction of: * = 0.05, ** = 0.01, *** = 0.001, **** = 0.0001, ns = not significant.

**Supplementary Figure 3.**
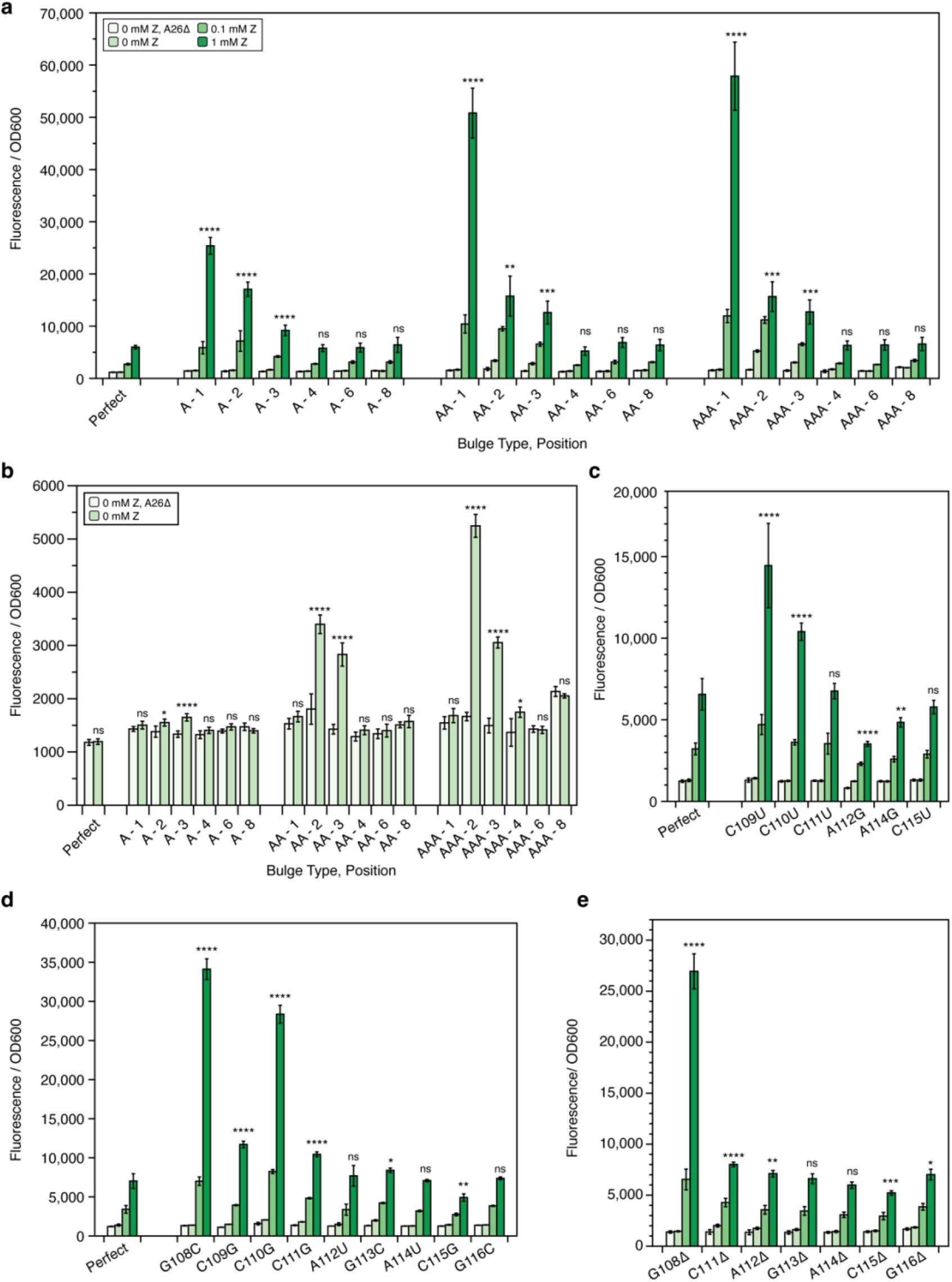
Fine-grained mutational analysis of invading strand reveals rules underlying control of dynamic range. (A) Bulge scans of the perfectly complementary invading strand demonstrate position-dependent effect of noncomplementary elements on riboswitch dynamic range. (B) A subset of the data from (A) is replotted to highlight differences in uninduced gene expression between the PK-incompetent background (A26Δ) and the PK-competent unmutated background. Larger kinetic barriers at positions 2 and 3 especially result in PK-dependent increases in leak. Mutational scans interrogating the effect of (C) wobble pairs, (D) mismatches, and (E) deletions demonstrate position-dependent effects of these noncomplementary elements on riboswitch dynamic range. Bars represent averages from three experimental replicates, each performed with triplicate biological replicates for a total of nine data points (n=9), with error bars representing standard deviation. T-test significance thresholds are for pairwise (two-tail, heteroscedastic) tests between ON-state (1 mM ZTP) FL/OD (A, C, D, E) or OFF-state (0 mM ZTP) FL/OD (B) with threshold values prior to Bonferroni correction of: * = 0.05, ** = 0.01, *** = 0.001, **** = 0.0001, ns = not significant.

**Supplementary Figure 4.**
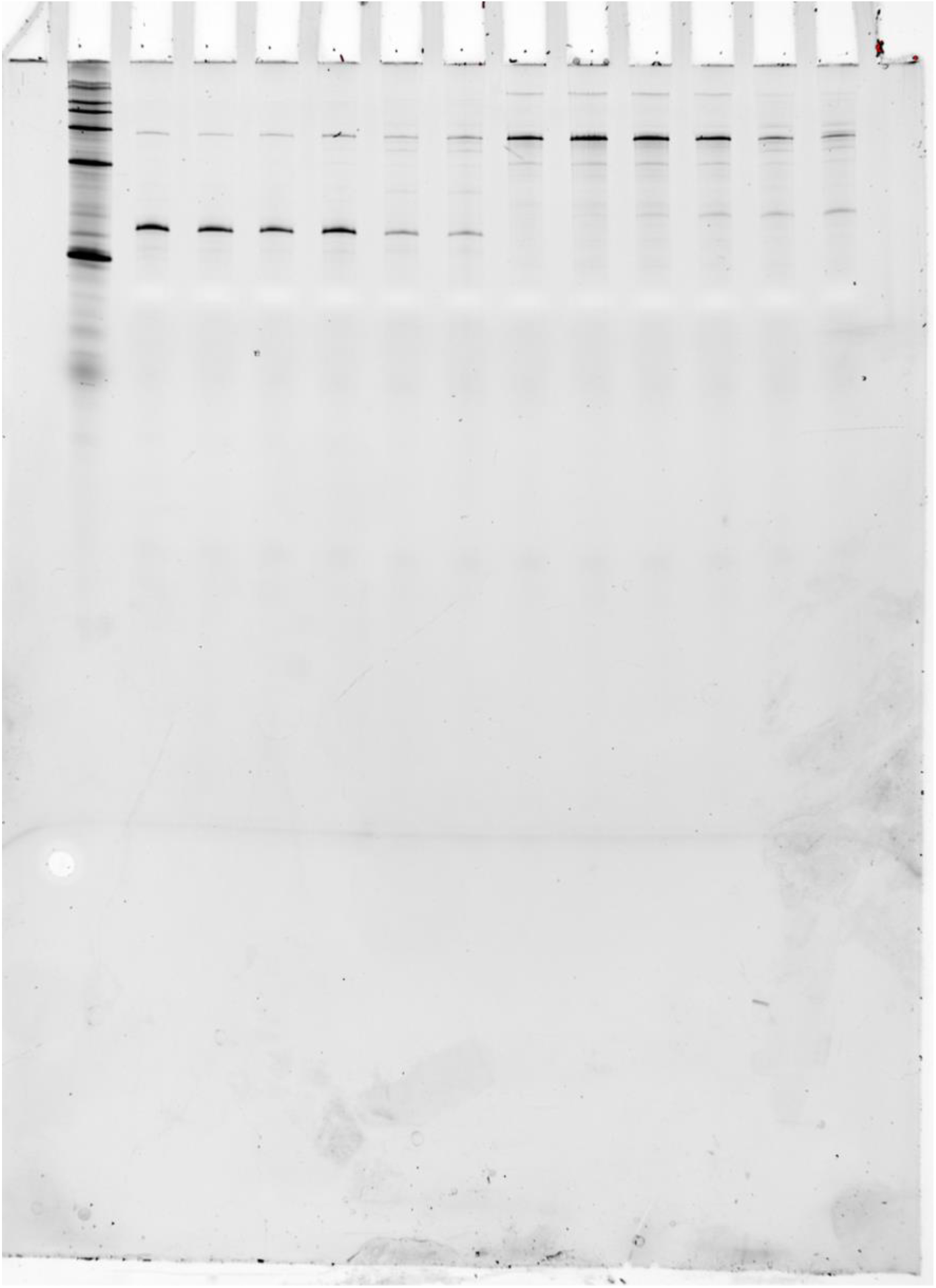
Figure 5B Source Gel 1. 1. RNA Century™-Plus Ladder (Invitrogen™) 2. Cbe wildtype, 0 µM Z, Replicate 1 3. Cbe wildtype, 10 µM Z, Replicate 1 4. Cbe wildtype, 32 µM Z, Replicate 1 5. Cbe wildtype, 100 µM Z, Replicate 1 6. Cbe wildtype, 320 µM Z, Replicate 1 7. Cbe wildtype, 1000 µM Z, Replicate 1 8. Cbe Flip, 0 µM Z, Replicate 1 9. Cbe Flip, 10 µM Z, Replicate 1 10. Cbe Flip, 32 µM Z, Replicate 1 11. Cbe Flip, 100 µM Z, Replicate 1 12. Cbe Flip, 320 µM Z, Replicate 1 13. Cbe Flip, 1000 µM Z, Replicate 1

**Supplementary Figure 5.**
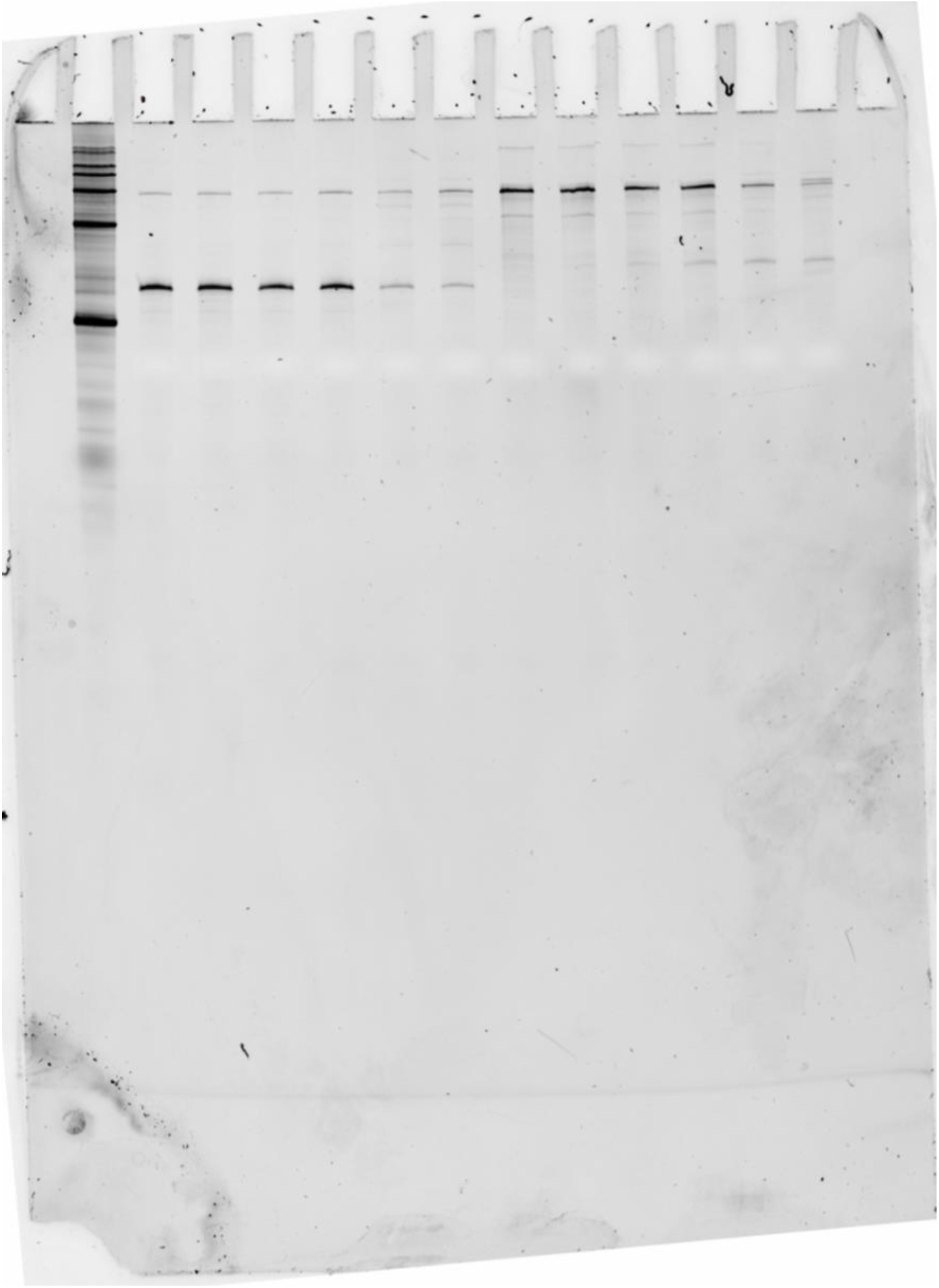
Figure 5B Source Gel 2. 1. RNA Century™-Plus Ladder (Invitrogen™) 2. Cbe wildtype, 0 µM Z, Replicate 2 3. Cbe wildtype, 10 µM Z, Replicate 2 4. Cbe wildtype, 32 µM Z, Replicate 2 5. Cbe wildtype, 100 µM Z, Replicate 2 6. Cbe wildtype, 320 µM Z, Replicate 2 7. Cbe wildtype, 1000 µM Z, Replicate 2 8. Cbe Flip, 0 µM Z, Replicate 2 9. Cbe Flip, 10 µM Z, Replicate 2 10. Cbe Flip, 32 µM Z, Replicate 2 11. Cbe Flip, 100 µM Z, Replicate 2 12. Cbe Flip, 320 µM Z, Replicate 2 13. Cbe Flip, 1000 µM Z, Replicate 2

**Supplementary Figure 6.**
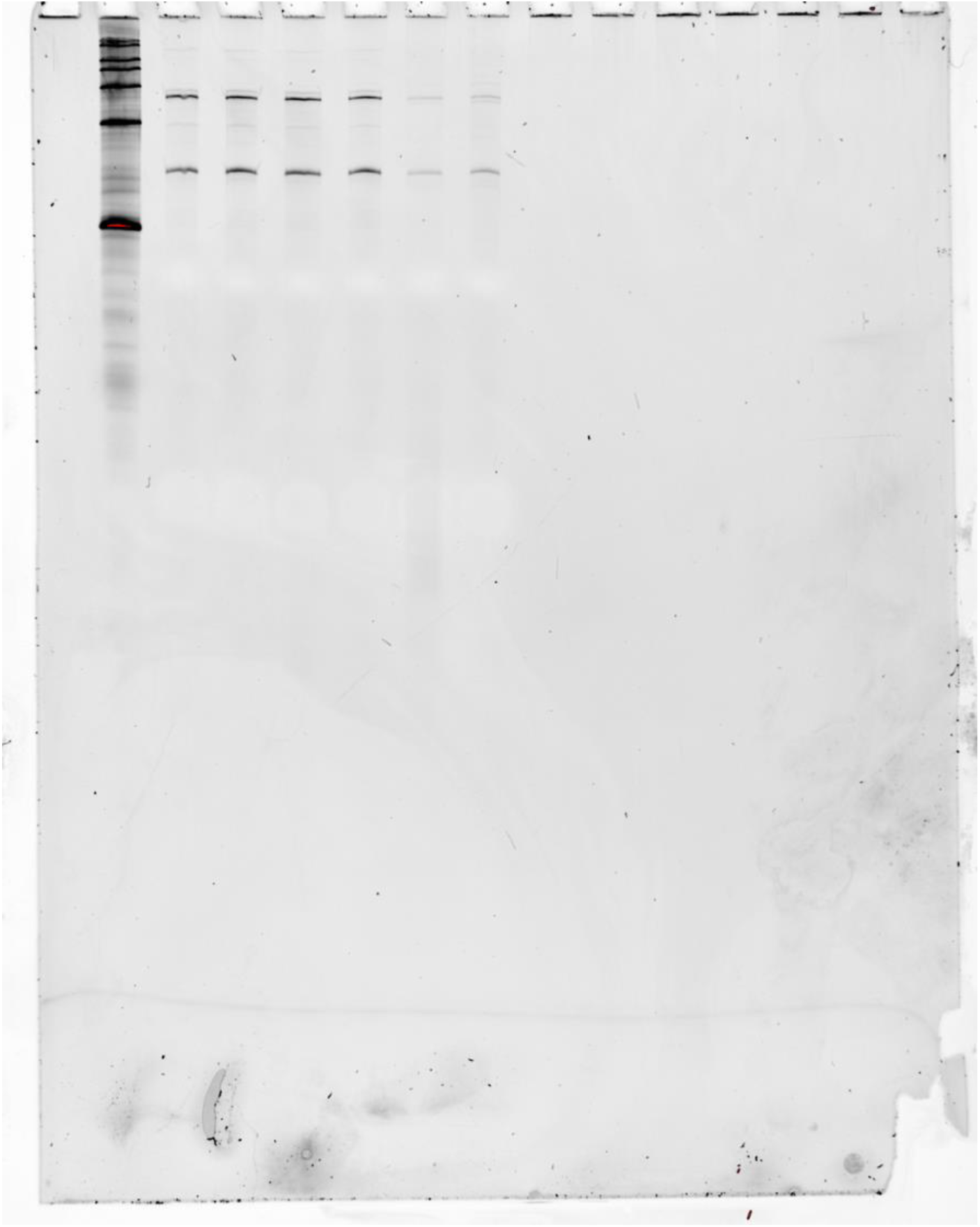
Figure 5B Source Gel 3. 1. RNA Century™-Plus Ladder (Invitrogen™) 2. Cbe AAA-Bulge Flip, 0 µM Z, Replicate 1 3. Cbe AAA-Bulge Flip, 10 µM Z, Replicate 1 4. Cbe AAA-Bulge Flip, 32 µM Z, Replicate 1 5. Cbe AAA-Bulge Flip, 100 µM Z, Replicate 1 6. Cbe AAA-Bulge Flip, 320 µM Z, Replicate 1 7. Cbe AAA-Bulge Flip, 1000 µM Z, Replicate 1

**Supplementary Figure 7.**
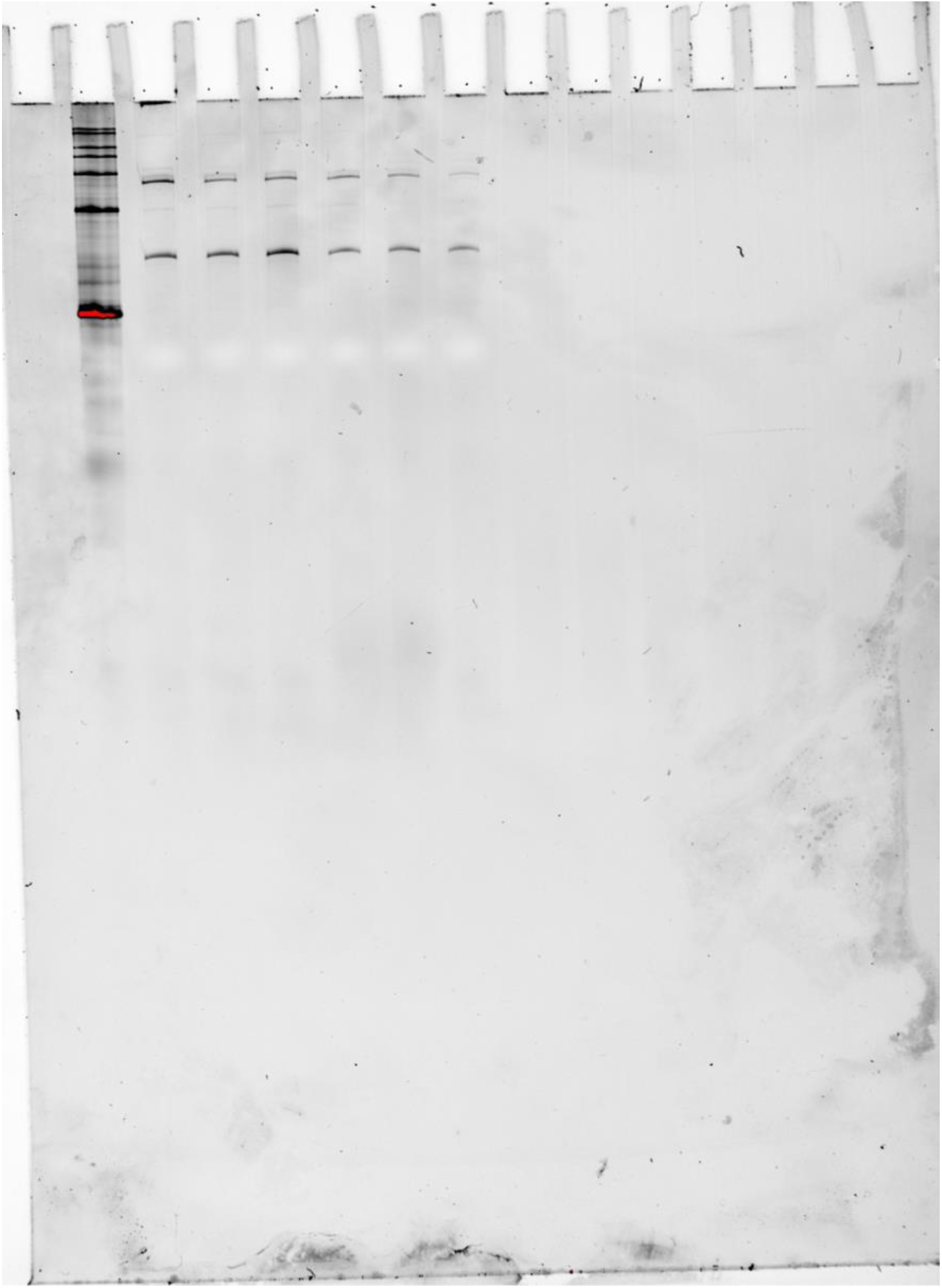
Figure 5B Source Gel 4. 1. RNA Century™-Plus Ladder (Invitrogen™) 2. Cbe AAA-Bulge Flip, 0 µM Z, Replicate 2 3. Cbe AAA-Bulge Flip, 10 µM Z, Replicate 2 4. Cbe AAA-Bulge Flip, 32 µM Z, Replicate 2 5. Cbe AAA-Bulge Flip, 100 µM Z, Replicate 2 6. Cbe AAA-Bulge Flip, 320 µM Z, Replicate 2 7. Cbe AAA-Bulge Flip, 1000 µM Z, Replicate 2

**Supplementary Table 1.**
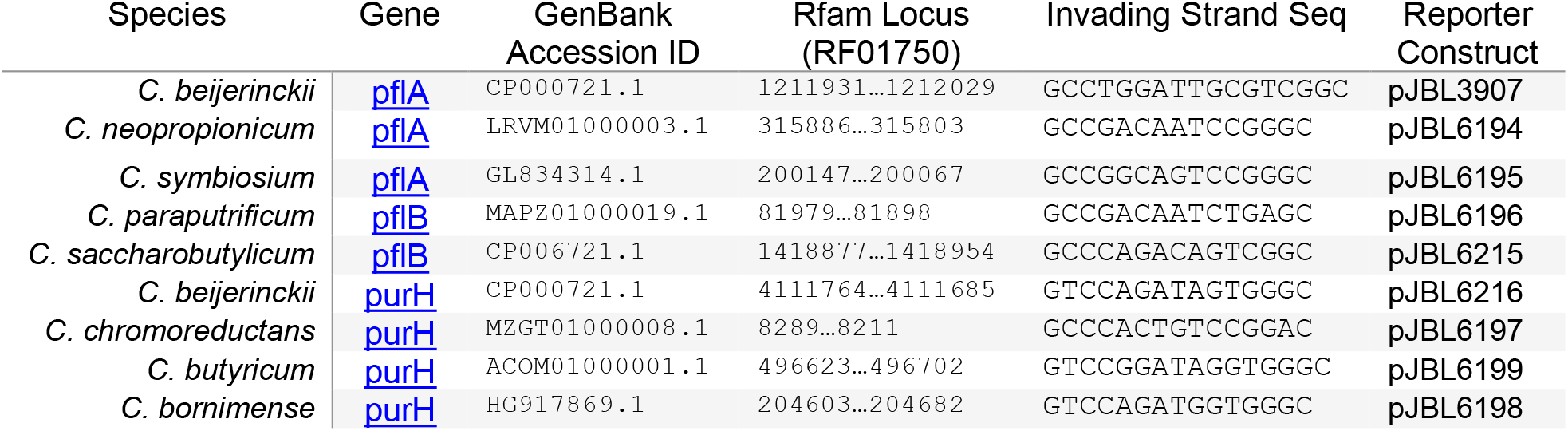
Source sequences used to generate chimeric riboswitches. For each invading strand sequence from the genus Clostridium that was used to generate a chimeric riboswitch, the species name, regulated gene (hyperlink to protein id), genome GenBank accession number, aptamer domain locus identified in Rfam, and invading strand sequence have been compiled. Sequences for indicated reporter constructs can be found in Supplemental Data File 1.

## Notes

### Competing Interest Statement

The authors have declared no competing interest.

